# Virulence phenotypes result from an interaction between pathogen ploidy and genetic background

**DOI:** 10.1101/672519

**Authors:** Dorian J. Feistel, Rema Elmostafa, Meleah A. Hickman

**Affiliations:** Department of Biology, Emory University

## Abstract

Studying fungal virulence is often challenging and frequently depends on many contexts, including host immune status and pathogen genetic background. However, ploidy has often been overlooked when studying virulence in eukaryotic pathogens. Since fungal pathogens, including the human opportunistic pathogen *Candida albicans*, can display extensive ploidy variation, assessing how ploidy impacts virulence has important clinical relevance. Here, we assessed how *C. albicans* ploidy and genetic background impact virulence phenotypes in both healthy and immunocompromised nematode hosts. In addition to reducing overall host survival, *Candida* negatively impacted host reproduction, which allowed us to survey lethal and non-lethal virulence phenotypes. While we did not detect any global differences in virulence between diploid and tetraploid pathogens, there were significant interactions between ploidy and *C. albicans* genetic background, regardless of host immune function.

## Introduction

Virulence is measured by the reduction of host fitness resulting from a host-pathogen interaction^1-3^. Therefore, virulence is not solely the property of the pathogen, but rather the product of the interaction between a host and its pathogen^4,5^. While many biotic and abiotic factors contribute to virulence^6,7^, the genotype-by-genotype interaction between hosts and pathogens is a primary determinant of whether a host gets infected and the resulting level of virulence^8,9^. One important, yet understudied, element of an organism’s genotype is its ploidy level. Theoretical studies demonstrate that host-pathogens co-evolution should drive hosts towards diploidy and pathogens towards haploidy^10^. However, polyploidy may have an important role in host-pathogen dynamics. For example, polyploidy of some host species is associated with elevated immune response^11,12^. Furthermore, polyploidy and aneuploidy are well documented in fungal pathogens that infect plant and/or animal hosts^13-15^. However there have only been a limited number of studies that investigate whether pathogen ploidy impacts virulence phenotypes and the few that have, often result in contradictory findings^13,16-19^.

One potential source of these contradictory results regarding ploidy is the genetic background or allelic composition of the pathogen. Allelic composition refers to not just the specific alleles present in a genome, but the amount of heterozygosity throughout the genome. Pathogen virulence depends on the pathogen’s specific allelic combination^20^ and phenotypic analysis of diverse clinical isolates within pathogenic species clearly demonstrate that pathogen genetic background contributes to its virulence^21,22^. Ploidy intrinsically impacts allelic composition – haploids contain a single set of gene alleles, whereas diploids and polyploids can either be homozygous or heterozygous for any given locus, and dominance can mask recessive alleles. Allelic composition in polyploids is further complicated by multiple allelic ratios, in which one to four alleles may be present, depending on the mechanism and age of the polyploidization event. Thus, a major challenge in determining the specific role ploidy has on pathogen virulence is disentangling it from allelic composition.

The opportunistic fungal pathogen *Candida albicans*, while typically a highly heterozygous diploid^23-25^, shows tremendous ploidy variation, ranging from haploid to polyploid^26-29^ and is highly tolerant of aneuploidy^26,30-34^. *C. albicans* is a commensal in the human gastrointestinal microbiota and various other niches^35^. Despite its commensal existence, *C. albicans* causes a range of infection, including superficial mucosal infections to life threatening systemic infections^36^. The severity of fungal infection is closely linked to the immune status of the host, with superficial mucosal infections occurring in healthy individuals and serious bloodstream infections occurring almost exclusively in immunocompromised hosts, the latter of which has a mortality of up to 50%^36^. Extensive allelic and ploidy variation, including loss-of-heterozygosity, aneuploidy and polyploidy, are well-documented in laboratory and clinical *C. albicans* isolates^21-25,37,38^. However, these large-scale genomic perturbations are often identified and studied in the context of antifungal drug resistance^22,31,32^, and only limited studies have specifically investigated how *C. albicans* ploidy impacts virulence phenotypes^16-18^.

In this study we sought to identify how *C. albicans* ploidy impacts its virulence by using four diploid-tetraploid pairs of strains, with each pair representing a distinct genetic background. We assessed virulence by monitoring four measures of host fitness using healthy and immunocompromised *C. elegans* hosts^39^. While we find almost no overall relationship between ploidy and virulence, there are detectable differences among *C. albicans* genetic backgrounds and clear interactions between *C. albicans* genetic background and its ploidy state on virulence phenotypes. We also observe these interactions in immunocompromised hosts, however, in some cases, the ploidy-specific pattern and/or the degree of virulence severity is different between healthy and immunocompromised hosts. Taken together, these results emphasize the importance of genotypic interactions on virulence phenotypes, in which genotype includes ploidy state.

## Results

### Interactions between pathogen ploidy and genetic background impact virulence phenotypes in healthy hosts

First, we wanted to investigate whether *C. albicans* genetic background differentially impacted host fitness. To do this we infected *C. elegans* hosts with four genetically distinct strain backgrounds of *C. albicans.* Two of these genetic backgrounds are laboratory strains: a ‘laboratory heterozygous,’ which consists of the SC5314 reference strain and a ‘laboratory homozygous,’ a derivative of SC5314 in which the genome is completely homozygous. The other two genetic backgrounds are clinical strains: a ‘bloodstream,’ isolated from a candidemia infection of a male immunosuppressed patient and an ‘oral/vaginal,’ a pair of clinical strains isolated from a single immune-competent female patient with vulvovaginal candidiasis. For each *C. albicans* genetic background, we assessed four measures of host fitness: host survival (Fig. 1A), host lineage growth (Fig. 1B), host brood size (Fig. 1C), and host reproductive timing (Fig. 1D). For survival and lineage growth, all *C. albicans* genetic backgrounds were virulent, with significant differences observed between uninfected and infected treatments, and the only statistical difference between *C. albicans* genetic backgrounds was between the laboratory heterozygous and oral/vaginal strains for host survival. For brood size, only the laboratory homozygous genetic background was virulent and resulted in significantly smaller host brood sizes compared to the two clinical *C. albicans* backgrounds. The impact on host reproductive timing also depended on pathogen genetic background: the laboratory heterozygous and the clinical bloodstream strains were virulent (i.e. reduced amount of normal reproductive timing) whereas the other two *C. albicans* genetic backgrounds were not. The laboratory heterozygous strain impacted host reproductive timing significantly more than the three other pathogen genetic backgrounds. Together, these results suggest that while pathogen genetic background may not obviously contribute to host survival phenotypes, it may be important for other measures of fungal infection that are less lethal.

**Figure 1:**
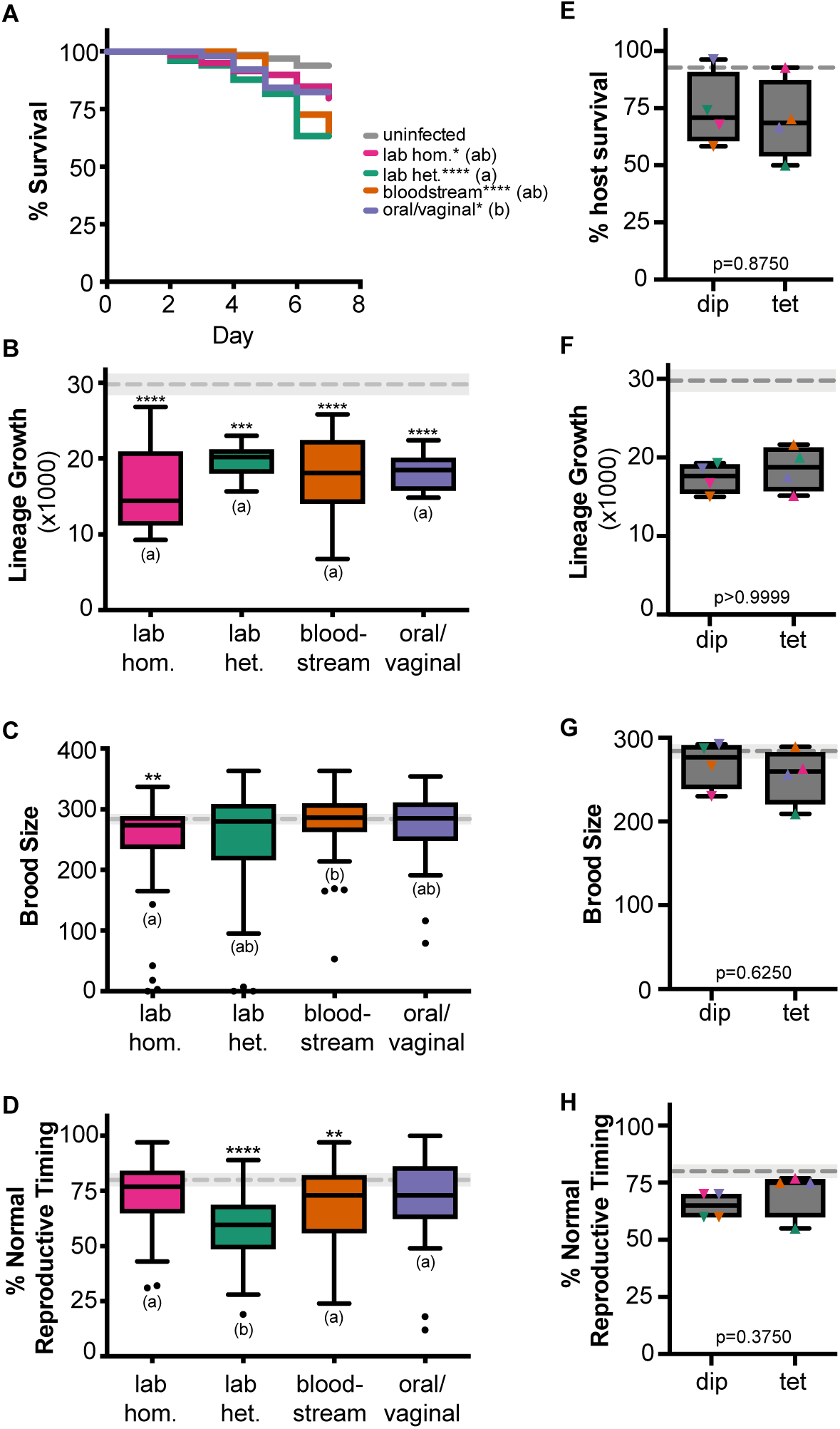
Different *C. albicans* genetic backgrounds reduce multiple measures of wildtype *C. elegans* host fitness. **A)** Survival curves for *C. elegans* populations that are either uninfected (exposed to just an E. coli food source, grey) or when infected with different *C. albicans* strains (indicated in legend). The number of worms analyzed (n) for each treatment is indicated in Table S1. Statistical significance was tested using pairwise comparisons of survival curves with Log-rank (Mantel-Cox) test. Astrisks denote statistical differences compared to the uninfected control (* indicates p <0.05, **** indicates p < 0.0001). *C. albicans* treatments that share letters are not significantly different, whereas treatments with differing letters are stastically different. **B)** Box and whiskers plot of host lineage growth which represents the total population size (representing the number of F1 and F2 progeny) produced within 7 days from a single founder *C. elegans* host infected with *C. albicans*. Boxes indicate the 25-75th quartiles with median indicated. Error bars are the normalized range of the data and circles indicate outliers. The mean and 95% CI of the uninfected control treatment are indicated by the grey dashed line and shaded grey box. Statistical significance was tested using one-way ANOVA. Astrisks denote statistical differences compared to the uninfected control (* indicates p <0.05, *** indicates p < 0.001). C. albicans treatments that share letters are not significantly different, whereas treatments with differing letters are stastically different, post-hoc Dunn’s multiple comparison test. **C)** Total brood size and **D)** Average percentage of host progeny produced during days 1-3 of adulthood (normal reproductive timing) of *C. elegans* infected with *C. albicans*. Data and statistical analysis are the same as (B). **E)** % host survival on Day 7 for diploid (dip) and tetraploid (tet) *C. albicans* strains (colored symbols indicate specific *C. albicans* genetic background). Statistical significance was tested using Wilcoxon matched-pairs signed rank test and p values are indicated. **F)** Lineage growth, **G)** Brood size, and **H)** Reproductive timing of hosts infected with *C. albicans* diploid and tetraploid strains. Data and statistical analysis are the same as for (E).

One factor that may dampen any differences in virulence among *C. albicans* genetic backgrounds is ploidy. For each genetic background, we used a related diploid strain and a tetraploid strain (Table S1). The tetraploids for both the ‘laboratory’ strains were produced via mating in the laboratory. The ‘bloodstream’ pair consisted of a diploid strain isolated early in the infection and its corresponding tetraploid strain was isolated mid-to late-infection following antifungal treatment. The ‘oral/vaginal’ pair consisted of a diploid strain isolated from the oral cavity and its corresponding tetraploid was isolated from the urogenital tract following antifungal treatment for vulvovaginal candidiasis. There have been limited and conflicted reports on the role of tetraploid in fungal virulence^16,17^. To assess how pathogen ploidy impacts virulence, we compared all the diploid strains to the tetraploid strains for host survival (Fig. 1E), lineage growth (Fig. 1F), host brood size (Fig. 1G), and host reproductive timing (Fig. 1H) in wildtype hosts. We did not observe a significant difference between these two ploidy states for any of the host fitness measures tested, suggesting that ploidy may not impact virulence phenotypes. However, when we account for *C. albicans* strain background, differences between pathogen ploidy emerge but depend on strain genetic background and thus, we detect a significant interaction between genetic background and ploidy for host lineage growth (‘interaction’ p=0.0354, two-way ANOVA; Fig. S1B), brood size (‘interaction’ p<0.0001, two-way ANOVA; Fig. S1B), and reproductive timing (‘interaction’ p=0.0283, two-way ANOVA; Fig. S1B). While we cannot directly test for an interaction using host survival curves, we do detect differences between diploids and tetraploids for each *C. albicans* genetic background (Fig. S1 and Table S2). Taken together, these results suggest that there is no global pattern in ploidy state and virulence, but ploidy in combination with genetic background does significantly contribute to *C. albicans* virulence phenotypes.

When we look at the specific diploid-tetraploid pairs, representing different *C. albicans* genetic backgrounds, we see significant differences in virulence between the diploid and tetraploid for at least one fitness measure, for every *C. albicans* genetic background (Fig. 2). For two genetic backgrounds, the laboratory homozygous and clinical bloodstream strains, the diploid strain was more virulent than its tetraploid counterpart. For the laboratory heterozygous and clinical oral/vaginal strains, the tetraploid strain was more virulent than its diploid counterpart, when differences between ploidy states were detected. Furthermore, these genetic background specific ploidy patterns are generally consistent across host fitness measures. For example, the laboratory heterozygous and clinical oral/vaginal tetraploids are also more virulent than their diploid counterparts for host brood size (Fig 2C), and the clinical bloodstream diploid was more virulent than its tetraploid counterpart for lineage growth and delayed host reproduction (Fig. 2B and D). We performed every pairwise comparison between treatments for host survival (Table S2), lineage growth (Table S3), brood size (Table S4), and reproductive timing (Table S5) and find significant differences in virulence between different *C. albicans* genetic backgrounds and ploidy states for most host fitness measures, except host lineage growth, were very few differences between *C. albicans* strains were detected. Taken together, these data support that *C. albicans* ploidy does contribute to its virulence phenotypes, but whether it attenuates or enhances virulence depends on its genetic background.

**Figure 2:**
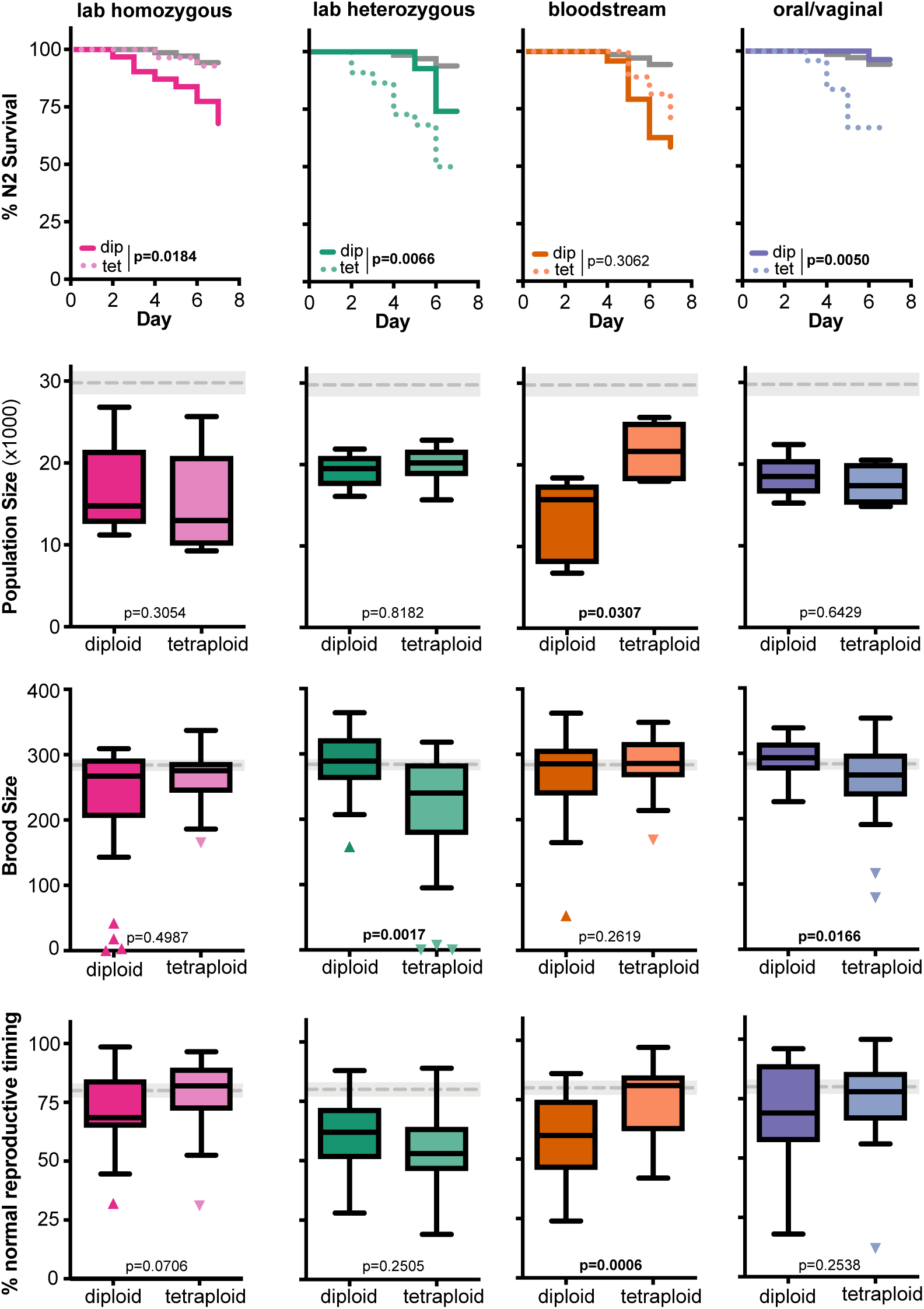
Ploidy-specific differences across *C. albicans* genetic backgrounds. **A)** Survival curves for *C. elegans* populations that are either uninfected (exposed to just an E. coli food source, grey) or when infected with diploid or tetraploid *C. albicans* strains from laboratory homozygous (pink), laboratory heterozygous (green), bloodstream (orange), or oral/vaginal (blue) genetic backgrounds. Statistical significance was tested using pairwise comparisons of diploid and tetraploid survival curves with Log-rank (Mantel-Cox) test and p-values are indicated and significant differences are highlighted in bold text. **B)** Box and whiskers plot of host lineage growth which represents the total population size (representing the number of F1 and F2 progeny) produced within 7 days from a single founder *C. elegans* host infected with *C. albicans*. Boxes indicate the 25-75th quartiles with median indicated. Error bars are the normalized range of the data and circles indicate outliers. The mean and 95% CI of the uninfected control treatment are indicated by the grey dashed line and shaded grey box. Statistical significance between diploid and tetraploid strains was determined using Mann Whitney test and p-values are indicated and significant differences are highlighted in bold text. **C)** Total brood size and **D)** Average percentage of host progeny produced during days 1-3 of adulthood (normal reproductive timing) of *C. elegans* infected with diploid and tetraploid *C. albicans*. Data and statistical analysis are the same as (B).

### Host immune status and pathogen genetic background contribute to virulence phenotypes

We have previously established that *C. albicans* and other non-*albicans Candida* species cause more severe infections in an immunocompromised *C. elegans* hosts compared to healthy *C. elegans* hosts ^39^. Given that we see pathogen ploidy patterns depend on genetic background for virulence phenotypes in wildtype healthy hosts, we next wanted to assess if we could detect comparable patterns in immunocompromised hosts. For each *C. albicans* strain, we assessed four measures of host fitness and compared *C. albicans* virulence in immunocompromised *sek-1 C. elegans* hosts: host survival (Fig. 3A), host lineage growth (Fig. 3B), host brood size (Fig. 3C), and host reproductive timing (Fig. 3D). Nearly all the *C. albicans* genetic backgrounds significantly reduced host fitness compared to uninfected controls, the only exception was the laboratory homozygous strains did not significantly delay host reproduction. The laboratory homozygous background is also less virulent than the clinical oral/vaginal genetic background for host. Together, these results suggest that global differences in pathogen genetic background are revealed in hosts with compromised immune function for both lethal and non-lethal measures of host fitness.

**Figure 3:**
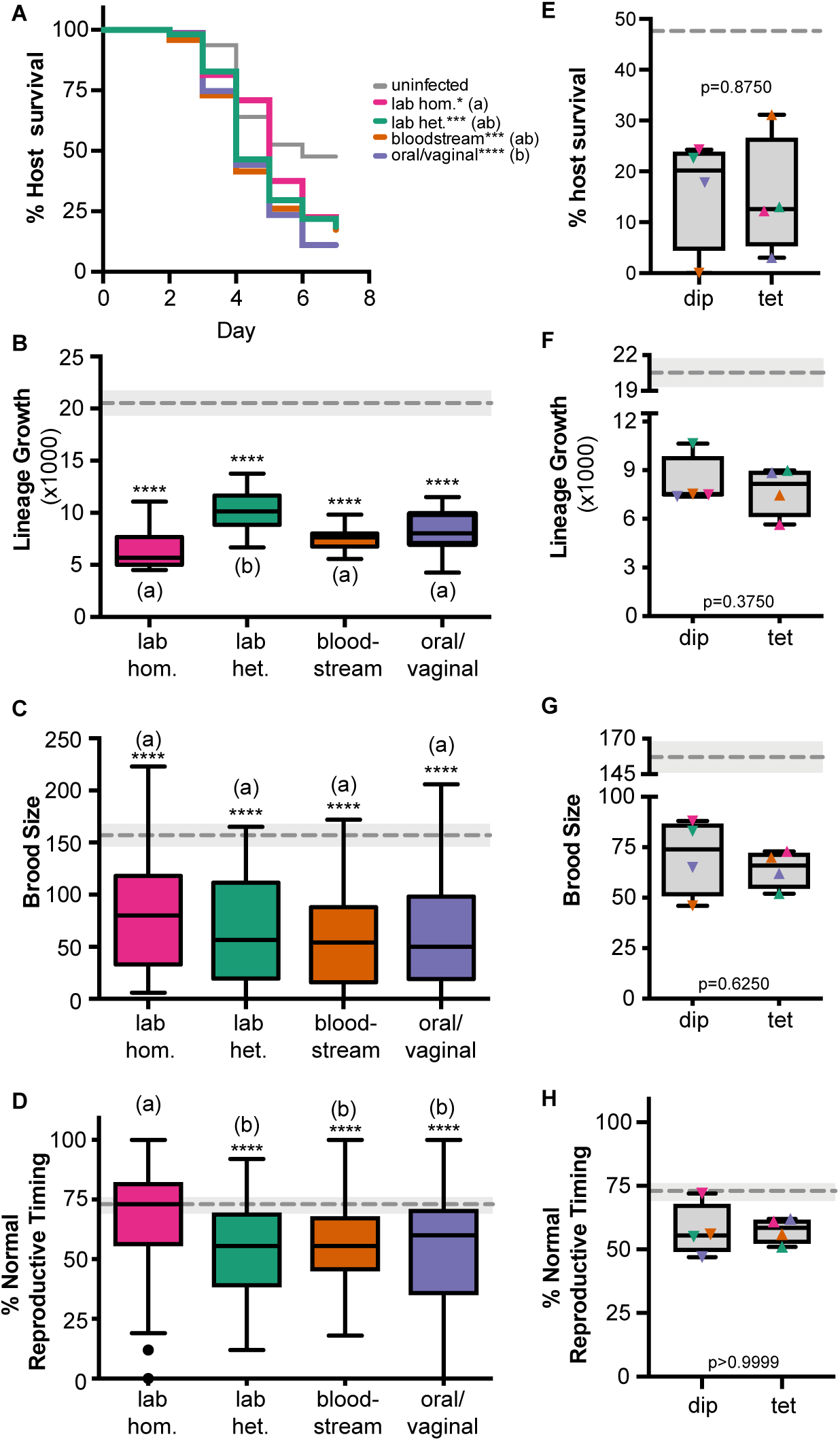
Immunocompromised hosts are highly susceptible to *C. albicans* infection regardless of genetic backgrounds or ploidy. **A)** Survival curves for *sek-1 C. elegans* populations that are either uninfected (exposed to just an *E. coli* food source, grey) or when infected with different *C. albicans* strains (indicated in legend). Statistical significance was tested using pairwise comparisons of survival curves with Log-rank (Mantel-Cox) test. Astrisks denote statistical differences compared to the uninfected control (* indicates p <0.05, **** indicates p < 0.0001). *C. albicans* treatments that share letters are not significantly different, whereas treatments with differing letters are stastically different. **B)** Box and whiskers plot of host lineage growth which represents the total population size (representing the number of F1 and F2 progeny) produced within 7 days from a single founder *sek-1 C. elegans* host infected with *C. albicans*. Boxes indicate the 25-75th quartiles with median indicated. Error bars are the normalized range of the data and circles indicate outliers. The mean and 95% CI of the uninfected control treatment are indicated by the grey dashed line and shaded grey box. Statistical significance was tested using one-way ANOVA. Astrisks denote statistical differences compared to the uninfected control (* indicates p <0.05, *** indicates p < 0.001). *C. albicans* treatments that share letters are not significantly different, whereas treatments with differing letters are stastically different, post-hoc Dunn’s multiple comparison test. **C)** Total brood size and **D)** Average percentage of host progeny produced during days 1-3 of adulthood (normal reproductive timing) of *sek-1 C. elegans* infected with *C. albicans*. Data and statistical analysis are the same as (B). **E)** % *sek-1* host survival on Day 7 for diploid (dip) and tetraploid (tet) *C. albicans* strains (colored symbols indicate specific *C. albicans* genetic background). Statistical significance between diploid and tetraploids was tested using Wilcoxon matched-pairs signed rank test and p values are indicated. **F)** Lineage growth, **G)** Brood size, and **H)** Reproductive timing of *sek-1* hosts infected with *C. albicans* diploid and tetraploid strains. Data and statistical analysis are the same as for (E).

To assess how pathogen ploidy impacts virulence in immunocompromised hosts, we compared all the diploid strains to the tetraploid strains for host survival (Fig. 3E), host lineage growth (Fig. 3F), host brood size (Fig. 3G), and host reproductive timing (Fig. 3H). Similar to healthy wildtype hosts, we did not observe a significant difference between these two ploidy states for and of the host fitness measures. However, when we account for *C. albicans* strain background, differences between pathogen ploidy emerge but depends on strain genetic background and a significant interaction between genetic background and ploidy is detected for host lineage growth (‘interaction’ p=0.0012, two-way ANOVA; Fig. S2B), brood size (‘interaction’ p=0.0090, two-way ANOVA; Fig. S2C), and reproductive timing (‘interaction’ p=0.0167, two-way ANOVA; Fig. S2D). While we cannot directly test for an interaction using host survival curves, we do detect differences between diploids and tetraploids for each *C. albicans* genetic background (Fig. S2 and Table S2). Taken together, these results further support the observation that there is no global correlation between ploidy state and virulence, but ploidy does have an important contribution to virulence phenotypes within pathogen genetic backgrounds regardless of host immune status.

When we look at specific diploid-tetraploid pairs, representing different *C. albicans* genetic background, we see significant differences in virulence in immunocompromised hosts between the diploid and tetraploid state for at least one fitness measure, for all *C. albicans* genetic backgrounds except for the laboratory homozygous (Fig. 4). For the laboratory heterozygous genetic background, the tetraploid counterpart was more virulent than its diploid counterpart when differences between ploidy states were detected (lineage growth and brood size), similar to the pattern observed in healthy hosts (Fig. 2). Furthermore, the clinical bloodstream diploid was more virulent than its tetraploid counterpart for immunocompromised host survival and brood size (Fig. 4A and C), similar to the pattern observed in healthy hosts. However, the oral/vaginal strain had significant differences between its diploid and tetraploid counterparts for host survival and reproductive timing (Fig. 4A and D), however the tetraploid was more virulent in the former, while the diploid was more virulent in the latter. We also performed every pairwise comparison between treatments for host survival (Table S2), lineage growth (Table S3), brood size (Table S4), and reproductive timing (Table S5) and find significant differences in virulence between different *C. albicans* genetic backgrounds and ploidy states for most host fitness measures, except host reproductive timing, were very few differences between *C. albicans* strains were detected. These results emphasize that while global differences between *C. albicans* genetic backgrounds or ploidy states may be difficult to identify in immunocompromised hosts (Fig. 3), ploidy-specific and genetic background differences in virulence are detectable across multiple host fitness measures.

**Figure 4:**
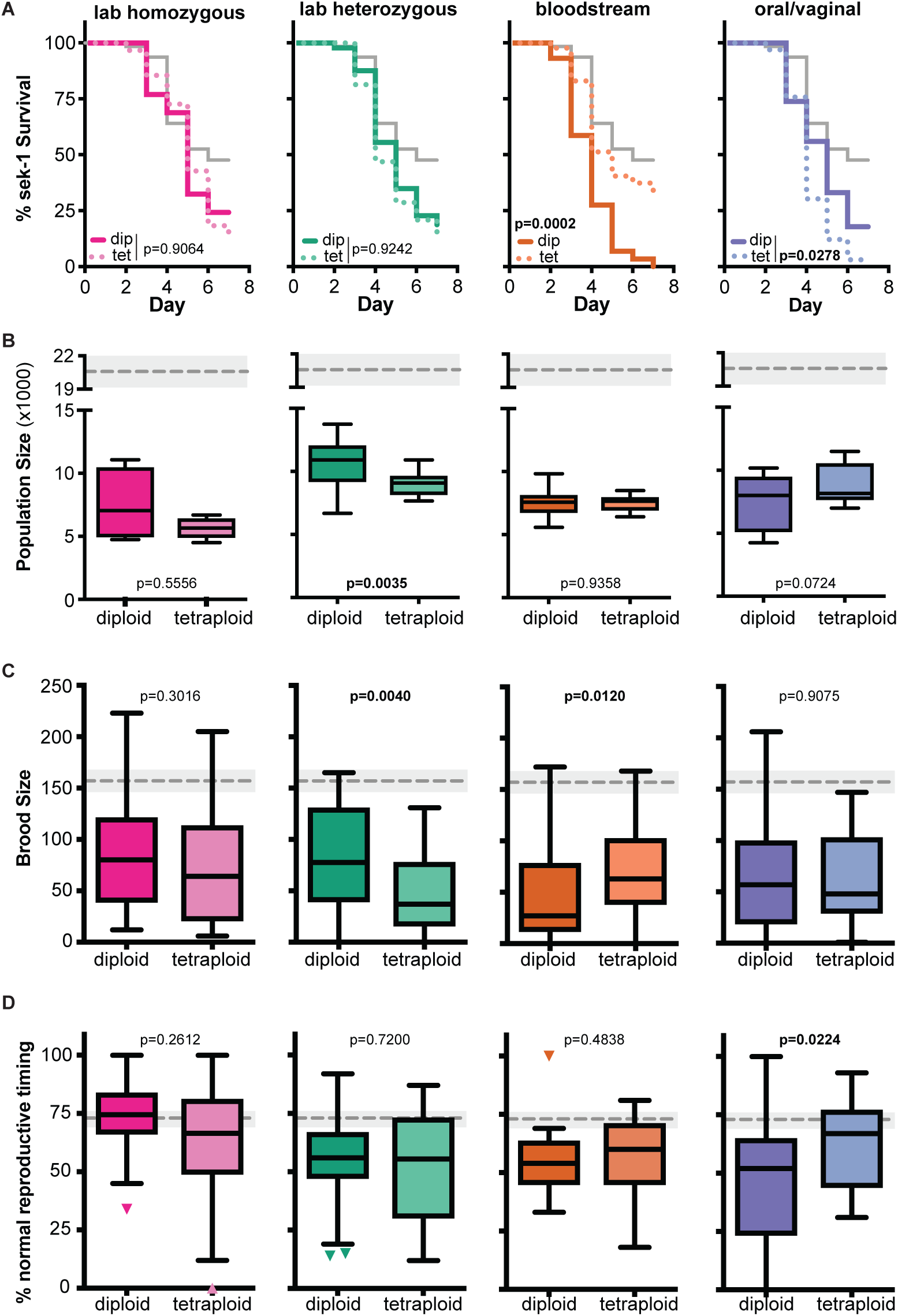
Ploidy-specific differences across *C. albicans* genetic backgrounds in immunocompromised hosts. **A)** Survival curves for *sek-1 C. elegans* populations that are either uninfected (exposed to just an E. coli food source, grey) or when infected with diploid or tetraploid *C. albicans* strains from laboratory homozygous (pink), laboratory heterozygous (green), bloodstream (orange), or oral/vaginal (blue) genetic backgrounds. Statistical significance was tested using pairwise comparisons of diploid and tetraploid survival curves with Log-rank (Mantel-Cox) test and p-values are indicated and significant differences are highlighted in bold text. **B)** Box and whiskers plot of host lineage growth which represents the total population size (representing the number of F1 and F2 progeny) produced within 7 days from a single founder *sek-1 C. elegans* host infected with *C. albicans*. Boxes indicate the 25-75th quartiles with median indicated. Error bars are the normalized range of the data and circles indicate outliers. The mean and 95% CI of the uninfected control treatment are indicated by the grey dashed line and shaded grey box. Statistical significance between diploid and tetraploid strains was determined using Mann Whitney test and p-values are indicated and significant differences are highlighted in bold text. **C)** Total brood size and **D)** Average percentage of host progeny produced during days 1-3 of adulthood (normal reproductive timing) of *sek-1 C. elegans* infected with diploid and tetraploid *C. albicans*. Data and statistical analysis are the same as (B).

### Ploidy-specific interactions between host and pathogen genotypes

Finally, we were curious if host immune status impacted the virulence relationship between each diploid-tetraploid pair of strains. Since there are significant differences for most host fitness measures between healthy and immunocompromised hosts even when uninfected (Table S6), we normalized each *C. albicans*-infected host fitness metric to that of the uninfected control to directly compare the severity of *C. albicans* infection between host genotypes. This analysis shows that immunocompromised hosts are more significantly more susceptible to *C. albicans* infection and show larger reductions in survival, brood size and lineage growth compared to those observed in healthy hosts, while the amount of delayed reproduction caused by *C. albicans* infection is similar (Fig. 5A-D). We also detect significant interactions between *C. albicans* strain and host immune status for lineage growth (‘interaction’ p=0.0004, two-way ANOVA), brood size (‘interaction’ p=0.0042, two-way ANOVA; Fig. S2C), and reproductive timing (‘interaction’ p=0.0001, two-way ANOVA).

**Figure 5:**
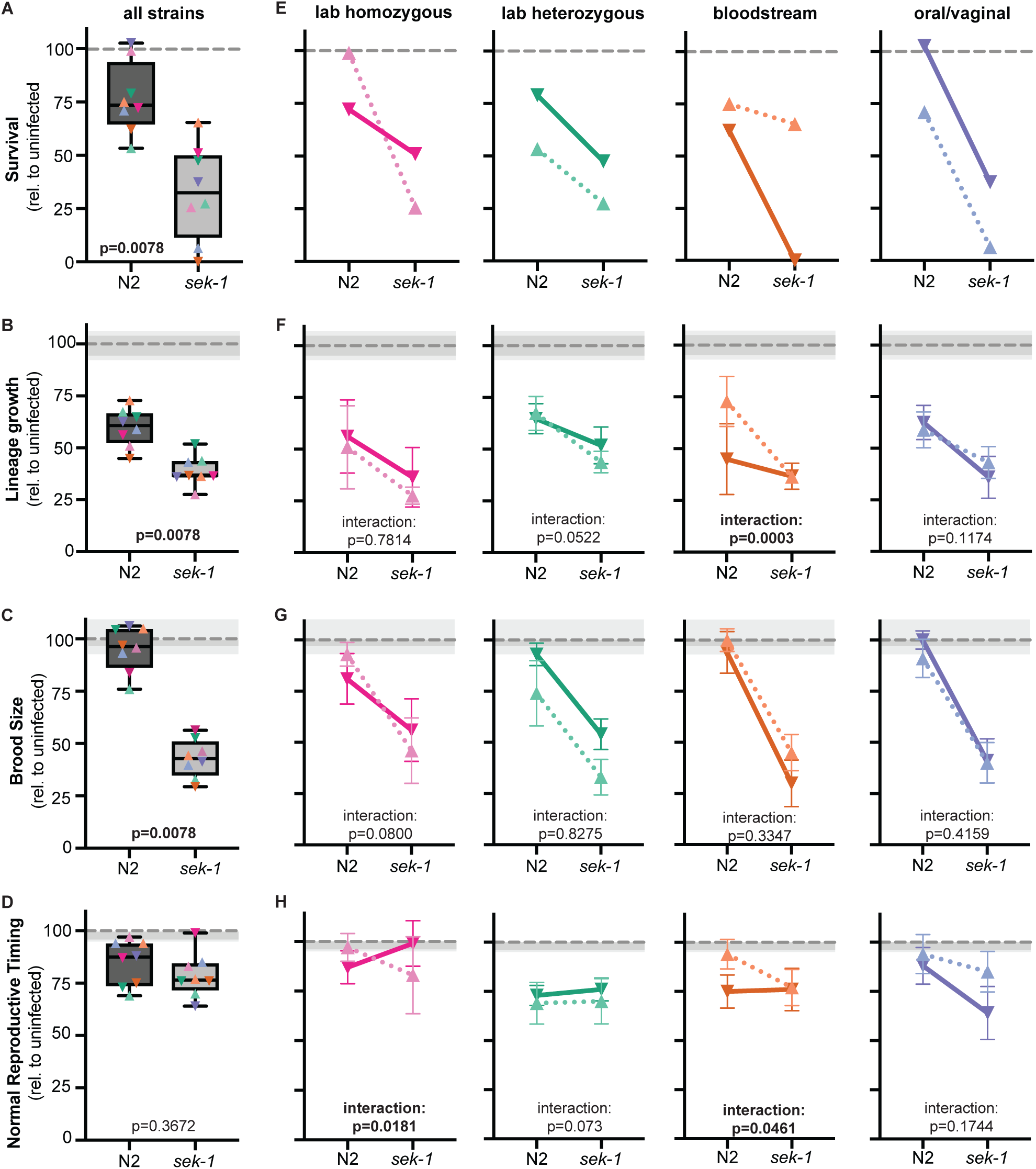
Ploidy-specific interactions between healthy and immunocompromised hosts. **A)** Relative impact on host survival from Day 7 survival data (*C. albicans* D7 survival/uninfected D7 survival for each host type) for all *C. albicans* strains (colored symbols indicate specific *C. albicans* genetic background) in healhty (N2) and immunocompromised hosts (*sek-1*). The mean and 95% CI of the uninfected control treatment are indicated by the grey dashed line and shaded box. Statistical significance between host genotypes was tested using Wilcoxon matched-pairs signed rank test and p values are indicated. **B)** Box and whiskers plot of relative host lineage growth, **C)** Brood size, and **D)** Reproductive timing between healthy. Data and statistical analysis are the same as (A). **E)** Relative impact of *C. albicans* ploidy on host survival. Relative virulence of diploid (solid lines) and tetraploid (dotted lines) *C. albicans* lab homozygous (pink), lab heterozygous (green), bloodstream (orange), and oral/vaginal (blue) genetic backgrounds in healthy (N2) and immunocompromised hosts (sek-1). Y-axis scale bar is the same as in (A). **F)** Relative host lineage growth, **G)** brood size, and **H)** reproductive timing between healthy and immunocompromised hosts across *C. albicans* genetic backgrounds. Symbols represent the mean value and error bars +/-SD. Y-axis scale bar is the same as in (B, C and D) respectively. Stastistical significance was tested by two-way ANOVA and ‘interaction’ p value indicated.

We next examined whether the relationship between ploidy-specific virulence differences changed depending on host immune status. To do this, we plotted the relative impact of the diploid (solid lines) and tetraploid (dotted lines) in healthy (N2) and immunocompromised (*sek-1*) host backgrounds for all four *C. albicans* genetic backgrounds (Fig 5E-H). In general, there was a high degree of similarity in the relationship between diploid and tetraploids across for both host genotypes, as indicated by a non-significant interaction term measured by two-way ANOVA. However, there were a couple of notable exceptions, particularly in the bloodstream pair of strains. In healthy hosts, we observed the diploid displaying more severe virulence phenotypes than its tetraploid counterpart for lineage growth and reproductive timing, yet these differences are diminished in immunocompromised hosts. Thus, we detect a significant interactions between host immune status and *C. albicans* ploidy for the bloodstream genetic background. Strikingly, we detect the reverse pattern for host survival with the *C. albicans* bloodstream diploid and tetraploid strains, in which no detectable differences are observed in healthy hosts, but the diploid is significantly more virulent than the tetraploid in immunocompromised hosts, yet regardless of the host context or specific fitness measure, the diploid is more virulent than its tetraploid counterpart. From these results, we conclude that interactions between host immune status, pathogen genetic background and ploidy state determine the severity of virulence phenotypes in *C. albicans*.

## Discussion

Here, we sought to understand how the genetic background and ploidy of the human fungal opportunistic pathogen *Candida albicans* contributes to virulence phenotypes in the nematode host *C. elegans*. Overall, we found that most of the strains we tested were virulent for at least one measure of host fitness, but cannot generalize any over-arching patterns in how ploidy or genetic background contribute to virulence phenotypes. Rather, we have found that there is a significant interaction between *C. albicans* ploidy and its genetic background. We detect this interaction in both healthy and immunocompromised host contexts. However, immunocompromised hosts display more severe fungal infections compared to healthy hosts, regardless of *C. albicans* ploidy and/or genetic background.

*C. albicans* was considered an ‘obligate’ heterozygous diploid for many years ^40-43^, and its diploid-tetraploid parasexual cycle only discovered and characterized in the last two decades^15,44^. Much of the research regarding *C. albicans* ploidy states have focused on the mechanisms involved with the parasexual cycle and how the ploidy reduction process that tetraploids undergo promote cellular heterogeneity and genetic variation^26,45-49^. There are only a small number of studies investigating if different ploidy states impact its virulence. These earlier studies investigating whether *C. albicans* ploidy impacts virulence had contradictory results – Hubbard et al demonstrated that derivative tetraploid had similar results in a murine tail vein infection model as its diploid progenitor^17^, whereas Ibrahim et al demonstrate that tetraploids are less virulent than diploids^16^. Given that tetraploid *C. albicans* have been identified in clinical settings^37^ and that other fungal pathogens display extensive ploidy variation^14,15^, assessing how ploidy impacts virulence has newfound clinical relevance. We think our systematic approach of comparing paired diploid and tetraploid strain across genetic backgrounds and host contexts provides a more comprehensive analysis of ploidy and virulence. Furthermore, our results support the contradictory findings reported earlier, as we observe that sometimes certain tetraploids are less virulent than diploids, sometimes there are no differences between ploidy states, and sometimes certain tetraploids are more virulent than diploids (Tables S2-S5).

Importantly, not only does our analysis account for differences in *C. albicans* genetic background, but, by utilizing *C. elegans* as an infection system we can assess multiple measures of host fitness beyond host survival. As an opportunistic pathogen, *C. albicans* causes a wide-range of infections with the most severe infections occurring in immunocompromised individuals and only a fraction of these severe infections result in patient death. As such, we need to broaden our scope to investigate fungal virulence beyond a lethal phenotype. We have previously demonstrated that *C. albicans* infection of *C. elegans* not only results in reduced survival, but also has reduced fecundity and delays in reproduction^39^. Here, we have leveraged this infection system to screen for differences in virulence between *C. albicans* ploidy and genetic backgrounds. While we did not find any overall pattern in virulence between diploids and tetraploids, we did identify ploidy-specific differences depending on genetic background (Figs. 2 and 4). Importantly, these specific patterns were consistent across multiple host fitness measures and host immune status (Fig. 5). Furthermore, we found that immunocompromised hosts had significantly more severe infections than immune competent hosts (Fig. 5), similar to what is observed clinically.

It is important to note that the clinical *C. albicans* genetic backgrounds, bloodstream and oral/vaginal, are not completely isogenic between the diploid and tetraploid strains, and that some allelic variation exists in addition to their differences in ploidy^37^. It is feasible that some of the differences in virulence that we observe between the diploid and tetraploids isolates in these two clinical *C. albicans* backgrounds (i.e. the bloodstream diploid is more virulent than its tetraploid, whereas the oral/vaginal tetraploid is more virulent than its diploid) is due to these allelic differences. By also measuring virulence phenotypes in laboratory-derived genomes that only differ in the number of chromosome sets they contain, we can directly assess the impact ploidy has on virulence. Here, we still observe different patterns of ploidy-specific virulence, depending on genetic background. In the laboratory heterozygous genome, we consistently observe the tetraploid as more virulent than its diploid in both healthy (Fig. 2) and immunocompromised (Fig. 4) host contexts. However, in the laboratory homozygous genome, we frequently failed to find any significant differences between diploid and tetraploid for any of the host fitness measures, the only exception being healthy host survival, in which the tetraploid was avirulent and the diploid was virulent (Fig. 2A). This result is consistent with previous findings that *C. albicans* homozygous genomes do not show significant growth differences *in vitro* or *in vivo* between ploidy states^27,46^.

In this work we found that ploidy and genetic background interact to contribute to *C. albicans* virulence (Figs. S1 and S2). We also observe significant interactions between *C. albicans* strains and host immune status (Fig. 5). These results indicate that virulence is not simply a binary ‘avirulent/virulent’ classification, but rather a complex trait and we need to start dissecting fungal virulence from this perspective. Recently, genome analysis of clinical isolates revealed genomic features that differ between clinical isolates and SC5314, the laboratory reference strain of *C. albicans*, and/or identify genetic determinants of antifungal drug resistance^22,38,50-53^. Only a small number have attempted to identify the genetic loci underpinning virulence and only two genes were identified and validated, *EFG1*^21,51^ and *PHO100*^46^. Importantly, these approaches may overlook other factors and contexts, such as ploidy or host immune status, that contribute to virulence. While it is clear that there variation in virulence occurs across *C. albicans* genetic backgrounds, there is still much work to do in elucidating the drivers of virulence.

## Material and Methods

### Strains and media

For this study, the *C. albicans* strains used are described in Table S1. *C. elegans* N2 Bristol (WT) and *sek-1* mutant^54^ were used to test host survival, fecundity and population growth in healthy and immunocompromised hosts, respectively. *C. elegans* populations were maintained at 20°C on lite nematode growth medium (NGM, US Biological) with *E. coli* OP50 as a food source and were transferred to a newly seeded *E. coli* plate every 3-4 days. For survival, fecundity and lineage growth assays, NGM was supplemented with 0.08 g/L uridine, 0.08 g/L histidine, and 0.04 g/L adenine to facilitate growth of auxotrophic *C. albicans* strains and 0.2 g/L streptomycin sulfate to inhibit *E. coli* overgrowth so *C. albicans* strains could proliferate.

### Survival and Fecundity Assays

Seeding NGM plates and *C. elegans* egg preparation for survival and fecundity assays were performed as previously described^39^. Briefly, *C. albicans* strains and *E. coli* OP50 were inoculated in 3 mL of YPD or 5 mL of LB, respectively, and incubated at 30°C for 1-2 days. *C. albicans* cultures were diluted to a final volume of 3.0 OD_600_ per mL and *E. coli* was pelleted and washed twice with 1 mL of ddH_2_O. The washed pellet was centrifuged for 60 sec at maximum and any excess liquid removed. The pellet was weighed and suspended with ddH_2_O to a final volume of 200 mg/mL. For uninfected treatments, 6.25 μL *E. coli* was mixed with 43.75 μL ddH_2_O. For *C. albicans* treatments, 1.25 μL *C. albicans* was mixed with 6.25 μL *E. coli* and brought to a final volume of 50 μL with ddH_2_O. All 50 μL was spotted onto the center of a 35mm supplemented-NGM Lite agar plate and incubated at room temperature overnight before addition of nematode eggs or transferred nematode.

To synchronize host populations, *C. elegans* populations (∼100 nematodes) were washed off NGM plates with M9 buffer and transferred to a 15 mL conical tube and pelleted by centrifugation for 2 min at 1200 rpm. The pellet was re-suspended in a 5.25% bleach solution, transferred to a microcentrifuge tube and inverted for 90-120 sec and subsequently centrifuged for 30 sec at 1500 rpm. The pellet was washed thrice with 1 mL M9 buffer and resuspended in 500 μL M9 buffer. ∼100 eggs were added to uninfected or *C. albicans* treatment plates. 48 h later, a single L4 nematode (x10 per treatment) was randomly selected and transferred to an uninfected or *C. albicans* treatment plate and incubated at 20°C. Each founder nematode was transferred to new treatment plates every 24 h for seven consecutive days. For survival analysis, we documented whether the founder was alive, dead, or censored (i.e. crawled off the plate or were otherwise unaccounted). For fecundity analysis, any eggs laid for each 24-hour interval remained undisturbed on the plate and incubated at 20°C for an additional 24 h and the number of viable progeny produced per day was scored. Total brood size is the sum of viable progeny produced over seven days. Delayed reproduction is calculated by the number of progeny produced on Day 4 or later divided by the total progeny produced for each founder nematode. All experiments were performed in triplicate or more.

### Lineage Growth Assays

Lineage growth assays were performed as previously described^39^. In brief, a single L4 founder nematode (x6 founders per treatment) was randomly selected from a synchronized population and transferred to a 100mm treatment plate that contained a 300 μl seed of *C. albicans* and/or *E. coli* and incubated at 20°C for five days in which the founder produces F1 and F2 progeny. The entire population derived from the single founder were washed with M9 buffer and transferred to 15mL conical tubes and brought to a final volume of 10mL. Tubes were placed at 4°C for 1h to allow the nematodes to settle at the bottom. 20 μL samples were taken from each population and counted six independent times to determine the final population size for each founder nematode. All experiments were performed at least in duplicate.

### Statistical Analyses

All statistical analyses were performed with GraphPad Prism. Survival curves were tested for significant differences using log-rank (Mantel Cox) tests. Lineage growth, total brood size and delayed reproduction data sets were tested for normality using the D’Agostino & Pearson omnibus normality test. For comparisons across genetic backgrounds, Kruskal-Wallis and Dunn’s multiple comparison tests were performed. For pairwise comparisons between ploidy or host context, Mann Whitney tests were performed. Two-way ANOVAs were performed to test for interactions between *C. albicans* genetic background and ploidy or between *C. albicans* ploidy and host context.

## Author contributions

DFJ and MAH devised the experiments. DFJ and RE performed the experiments. DJF and MAH analyzed the data. MAH wrote the manuscript.

