## Supplemental Files for "Virulence phenotypes result from an interaction between pathogen ploidy and genetic background"

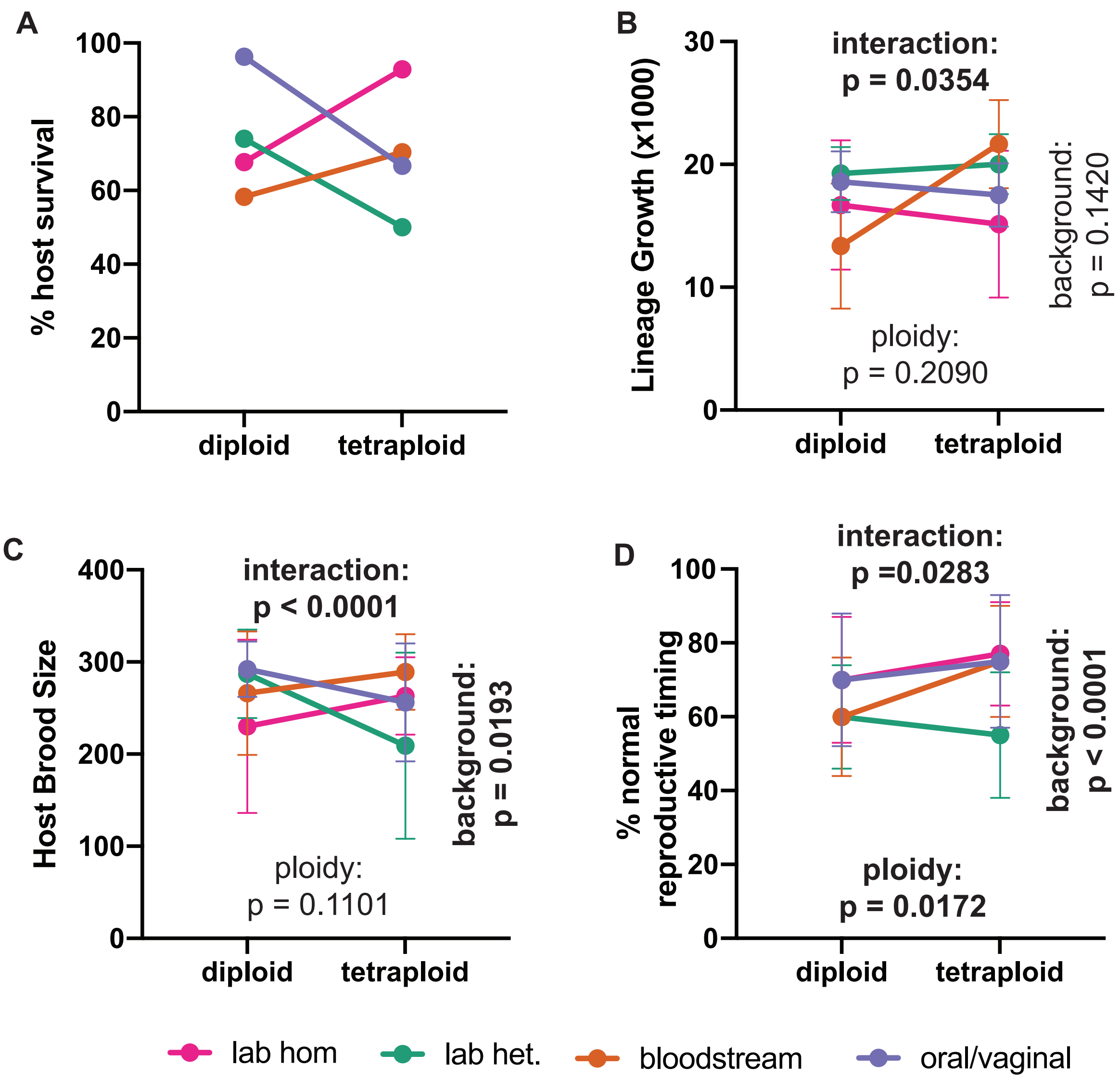


**Figure S1: Interaction between C. albicans ploidy and genetic background on virulence phenotypes in healthy hosts.**

**A)** Relationship of Day 7 host survival between diploid or tetraploid *C. albicans* of the lab homozygous (pink), lab heterozygous (green), bloodstream (orange), and oral/vaginal (blue) genetic backgrounds in healthy (N2) hosts.

**B)** Relationship of host lineage growth, **C)** brood size, and **D)** reproductive timing between diploid or tetraploid *C. albicans* of the lab homozygous (pink), lab heterozygous (green), bloodstream (orange), and oral/vaginal (blue) genetic backgrounds in healthy (N2) hosts. Symbols represent the mean value and error bars +/-SD. Stastistical significance was tested by two-way ANOVA and p values for 'ploidy,' 'genetic background,' and their 'interaction' is indicated.

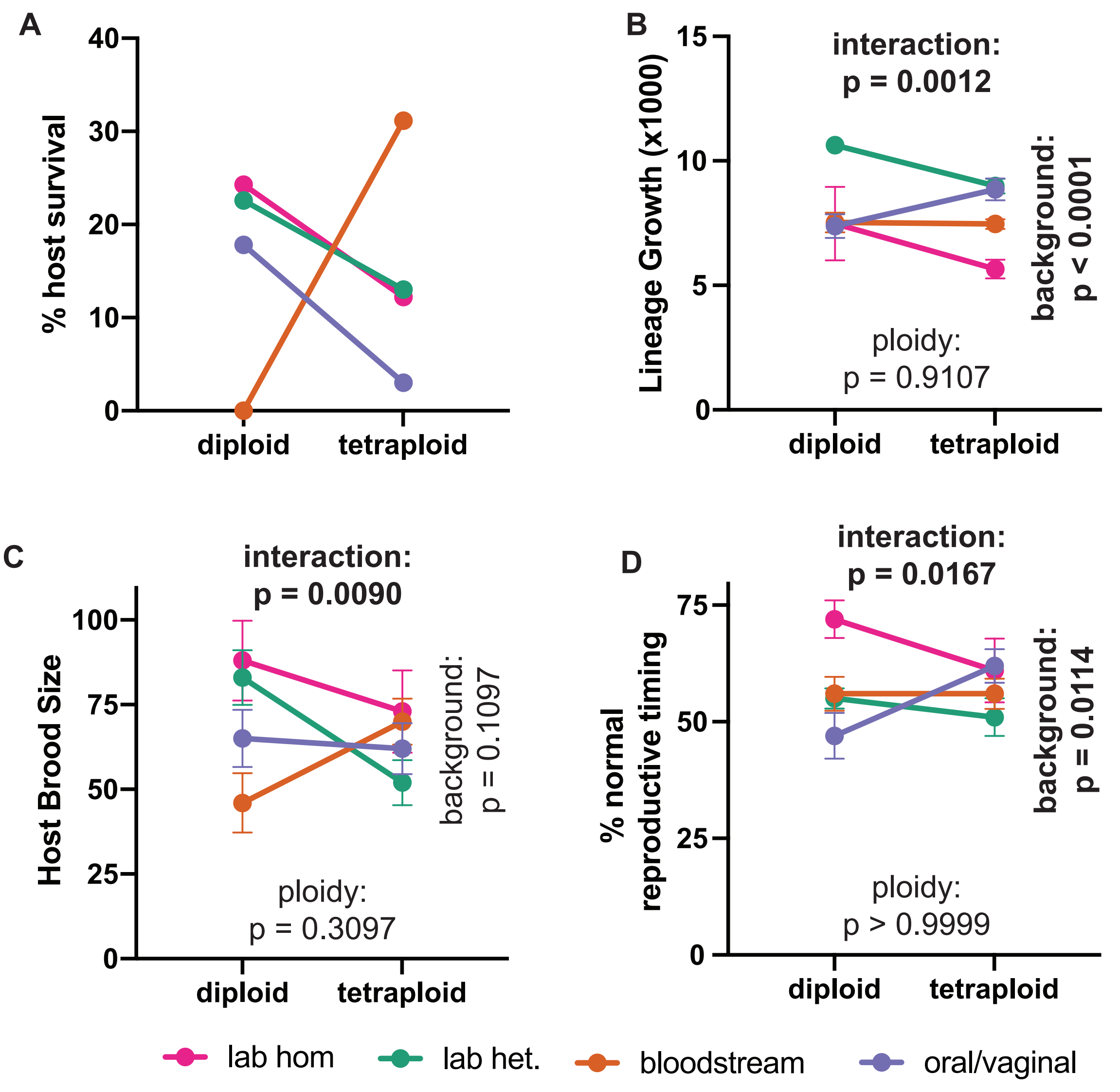


**Figure S2: Interaction between *C. albicans* ploidy and genetic background on virulence phenotypes in immunocompromised hosts.**

**A)** Relationship of Day 7 host survival between diploid or tetraploid *C. albicans* of the lab homozygous (pink), lab heterozygous (green), bloodstream (orange), and oral/vaginal (blue) genetic backgrounds in immunocompromised (*sek-1*) hosts.

**B)** Relationship of host lineage growth, **C)** brood size, and **D)** reproductive timing between diploid or tetraploid *C. albicans* of the lab homozygous (pink), lab heterozygous (green), bloodstream (orange), and oral/vaginal (blue) genetic backgrounds in immunocompromised (*sek-1*) hosts. Symbols represent the mean value and error bars +/-SD. Stastistical significance was tested by two-way ANOVA and p values for 'ploidy,' 'genetic background,' and their 'interaction' is indicated.

| Strain | Alias | Ploidy | Genetic Background | Source |
|---|---|---|---|---|
| YJB12804 | lab hom. | diploid | Laboratory SC5314-derived homozygous genome (auto-diploid of YJBXXX) | Hickman et al 2013; Gerstein et al (2017) |
| MH306 | lab hom. | tetraploid | Mating product between two SC5314-derived homozygous strains | This study |
| SC5314 | lab het. | diploid | Laboratory reference strain | Gillum et al 1984 |
| RBY18 | lab het. | tetraploid | Mating product between two SC5314-derived heterozygous strains | Bennett and Johnson 2003 |
| FH1 | bloodstream | diploid | Clinical isolate from marrow transplant patient | Hull et al Marr et al (1997); Abbey et al (2014) |
| FH6 | bloodstream | ~tetraploid | Clinical isolate from marrow transplant patient (same patient as FH1) | Hull et al Marr et al (1997); Abbey et al (2014) |
| PN2 | oral/vaginal | diploid | Clinical isolate recovered from oral cavity | Gerstein et al (2017) |
| PN1 | oral/vaginal | tetraploid | Clinical isolate recovered from vaginal infection (same patient as PN2) | Gerstein et al (2017) |

Table S1. Strains used in this study

| Strain 1 | Strain 2 | Healthy hosts | Immunocompromised hosts |
|---|---|---|---|
| | | p-value | p-value |
| uninfected | 2C Lab hom | ***** (0.0003)** | ns (0.0679) |
| | 4C Lab hom | ns (0.7882) | *** (0.0484)** |
| | 2C Lab het | ****** (<0.0001)** | **** (0.0017)** |
| | 4C lab het | ****** (<0.0001)** | ***** (0.0005)** |
| | 2C bloodstream | ****** (<0.0001)** | ****** (<0.0001)** |
| | 4C bloodstream | **** (0.0015)** | ns (0.0777) |
| | 2C oral/vag | ns (0.6802) | **** (0.0029)** |
| | 4C oral/vag | ***** (0.0002)** | ****** (<0.0001)** |
| 2C Lab hom | 4C Lab hom | *** (0.0184)** | ns (0.9065) |
| | 2C Lab het | *** (0.0175)** | ns (0.6837) |
| | 4C lab het | ***** (0.0002)** | ns (0.2758) |
| | 2C bloodstream | ns (0.5167) | ***** (0.0006)** |
| | 4C bloodstream | ns (0.7351) | ns (0.8526) |
| | 2C oral/vag | **** (0.0060)** | ns (0.5026) |
| | 4C oral/vag | ns (0.8687) | **** (0.0076)** |
| 4C Lab hom | 2C Lab het | ****** (<0.0001)** | ns (0.8114) |
| | 4C lab het | ****** (<0.0001)** | ns (0.3901) |
| | 2C bloodstream | **** (0.0038)** | ***** (0.0004)** |
| | 4C bloodstream | *** (0.0376)** | ns (0.6867) |
| | 2C oral/vag | ns (0.5683) | ns (0.5807) |
| | 4C oral/vag | *** (0.0164)** | **** (0.0050)** |
| 2C lab het | 4C lab het | **** (0.0066)** | ns (0.9242) |
| | 2C bloodstream | ns (0.0985) | ****** (<0.0001)** |
| | 4C bloodstream | **** (0.0061)** | ns (0.4505) |
| | 2C oral/vag | ****** (<0.0001)** | ns (0.6232) |
| | 4C oral/vag | ns (0.0620) | **** (0.0019)** |
| 4C lab het | 2C bloodstream | **** (0.0019)** | **** (0.0043)** |
| | 4C bloodstream | ****** (<0.0001)** | ns (0.1528) |
| | 2C oral/vag | ****** (<0.0001)** | ns (0.7079) |
| | 4C oral/vag | **** (0.0014)** | *** (0.0477)** |
| 2C bloodstream | 4C bloodstream | ns (0.3062) | ***** (0.0002)** |
| | 2C oral/vag | ***** (0.0009)** | **** (0.0021)** |
| | 4C oral/vag | ns (0.7936) | ns (0.2983) |
| 4C bloodstream | 2C oral/vag | *** (0.0114)** | ns (0.3112) |
| | 4C oral/vag | ns (0.5600) | **** (0.0029)** |
| 2C oral/vag | 4C oral/vag | **** (0.0050)** | *** (0.0278)** |

Table S2: Pairwise survival curve comparisons (log-rank test) for uninfected and all eight *C. albicans* strains, for each host genotype.

| Strain 1 | Strain 2 | Healthy hosts | Immunocompromised hosts |
|---|---|---|---|
| | | p-value | p-value |
| uninfected | 2C Lab hom | ****** (<0.0001)** | ****** (<0.0001)** |
| | 4C Lab hom | ****** (<0.0001)** | ****** (<0.0001)** |
| | 2C Lab het | ****** (<0.0001)** | ****** (<0.0001)** |
| | 4C lab het | ****** (<0.0001)** | ****** (<0.0001)** |
| | 2C bloodstream | ****** (<0.0001)** | ****** (<0.0001)** |
| | 4C bloodstream | ***** (0.0006)** | ****** (<0.0001)** |
| | 2C oral/vag | ****** (<0.0001)** | ****** (<0.0001)** |
| | 4C oral/vag | ***** (0.0003)** | ****** (<0.0001)** |
| 2C Lab hom | 4C Lab hom | ns (0.3054) | ns (0.5556) |
| | 2C Lab het | ns (0.2198) | *** (0.0282)** |
| | 4C lab het | ns (0.1471) | ns (0.4605) |
| | 2C bloodstream | ns (0.5135) | ns (0.8513) |
| | 4C bloodstream | ns (0.0753) | ns (>0.9999) |
| | 2C oral/vag | ns (0.2635) | ns (0.8313) |
| | 4C oral/vag | ns (0.5395) | ns (0.4626) |
| 4C Lab hom | 2C Lab het | ns (0.2520) | ****** (<0.0001)** |
| | 4C lab het | ns (0.1120) | ***** (0.0002)** |
| | 2C bloodstream | ns (0.4222) | **** (0.0087)** |
| | 4C bloodstream | ns (0.0709) | ***** (0.0009)** |
| | 2C oral/vag | ns (0.3002) | ns (0.1341) |
| | 4C oral/vag | ns (0.5185) | ***** (0.0002)** |
| 2C lab het | 4C lab het | ns (0.8182) | **** (0.0035)** |
| | 2C bloodstream | *** (0.0303)** | ****** (<0.0001)** |
| | 4C bloodstream | ns (0.3290) | ****** (<0.0001)** |
| | 2C oral/vag | ns (0.6991) | ****** (<0.0001)** |
| | 4C oral/vag | ns (0.2857) | ns (0.2439) |
| 4C lab het | 2C bloodstream | *** (0.0303)** | **** (0.0069)** |
| | 4C bloodstream | ns (0.5368) | ***** (0.0004)** |
| | 2C oral/vag | ns (0.2403) | *** (0.0396)** |
| | 4C oral/vag | ns (0.2571) | ns (0.5698) |
| 2C bloodstream | 4C bloodstream | *** (0.0317)** | ns (0.9358) |
| | 2C oral/vag | ns (0.0823) | ns (0.9241) |
| | 4C oral/vag | ns (0.2857) | *** (0.0490)** |
| 4C bloodstream | 2C oral/vag | ns (0.3290) | ns (0.8240) |
| | 4C oral/vag | ns (0.1905) | *** (0.0147)** |
| 2C oral/vag | 4C oral/vag | ns (0.6429) | ns (0.0724) |

Table S3: Pairwise lineage growth comparisons (Mann Whitney test) for uninfected and all eight *C. albicans* strains, for each host genotype.

| Strain 1 | Strain 2 | Healthy hosts | Immunocompromised hosts |
|---|---|---|---|
| | | p-value | p-value |
| uninfected | 2C Lab hom | **** (0.0023)** | ****** (<0.0001)** |
| | 4C Lab hom | *** (0.0246)** | ****** (<0.0001)** |
| | 2C Lab het | ns (0.4481) | ****** (<0.0001)** |
| | 4C lab het | ***** (0.0004)** | ****** (<0.0001)** |
| | 2C bloodstream | ns (0.5219) | ****** (<0.0001)** |
| | 4C bloodstream | ns (0.5210) | ****** (<0.0001)** |
| | 2C oral/vag | ns (0.2424) | ****** (<0.0001)** |
| | 4C oral/vag | ns (0.0511) | ****** (<0.0001)** |
| 2C Lab hom | 4C Lab hom | ns (0.4987) | ns (0.3016) |
| | 2C Lab het | **** (0.0068)** | ns (0.7813) |
| | 4C lab het | ns (0.3550) | *** (0.0120)** |
| | 2C bloodstream | ns (0.1106) | **** (0.0038)** |
| | 4C bloodstream | **** (0.0027)** | ns (0.2498) |
| | 2C oral/vag | **** (0.0012)** | ns (0.1265) |
| | 4C oral/vag | ns (0.4865) | ns (0.1646) |
| 4C Lab hom | 2C Lab het | *** (0.0257)** | ns (0.4235) |
| | 4C lab het | ns (0.0637) | ns (0.1784) |
| | 2C bloodstream | ns (0.3451) | *** (0.0500)** |
| | 4C bloodstream | *** (0.0314)** | ns (0.9242) |
| | 2C oral/vag | ***** (0.0061)** | ns (0.5840) |
| | 4C oral/vag | ns (0.9819) | ns (0.7950) |
| 2C lab het | 4C lab het | **** (0.0017)** | ***** (0.0010)** |
| | 2C bloodstream | ns (0.3270) | **** (0.0021)** |
| | 4C bloodstream | ns (0.9897) | ns (0.2655) |
| | 2C oral/vag | ns (0.7946) | ns (0.0817) |
| | 4C oral/vag | ns (0.0620) | ns (0.0746) |
| 4C lab het | 2C bloodstream | *** (0.0353)** | ns (0.2879) |
| | 4C bloodstream | **** (0.0012)** | *** (0.0364)** |
| | 2C oral/vag | ***** (0.0002)** | ns (0.3985) |
| | 4C oral/vag | ns (0.1231) | ns (0.2828) |
| 2C bloodstream | 4C bloodstream | ns (0.2619) | *** (0.0102)** |
| | 2C oral/vag | ns (0.1678) | ns (0.1165) |
| | 4C oral/vag | ns (0.4100) | *** (0.0497)** |
| 4C bloodstream | 2C oral/vag | ns (0.8151) | ns (0.4683) |
| | 4C oral/vag | *** (0.0393)** | ns (0.3723) |
| 2C oral/vag | 4C oral/vag | *** (0.0166)** | ns (0.9075) |

Table S4: Pairwise brood size comparisons (Mann Whitney test) for uninfected and all eight *C. albicans* strains, for each host genotype.

| Strain 1 | Strain 2 | Healthy hosts (N2) | Immunocompromised hosts (sek-1) |
|---|---|---|---|
| | | p-value | p-value |
| uninfected | 2C Lab hom | **** (0.0062)** | ns (0.8702) |
| | 4C Lab hom | ns (0.4611) | ns (0.1466) |
| | 2C Lab het | ****** (<0.0001)** | ****** (<0.0001)** |
| | 4C lab het | ****** (<0.0001)** | ****** (<0.0001)** |
| | 2C bloodstream | ****** (<0.0001)** | ****** (<0.0001)** |
| | 4C bloodstream | ns (0.2276) | ****** (<0.0001)** |
| | 2C oral/vag | *** (0.0235)** | ****** (<0.0001)** |
| | 4C oral/vag | ns (0.2811) | **** (0.0065)** |
| 2C Lab hom | 4C Lab hom | ns (0.0706) | ns (0.2612) |
| | 2C Lab het | *** (0.0147)** | ***** (0.0002)** |
| | 4C lab het | **** (0.0051)** | **** (0.0017)** |
| | 2C bloodstream | *** (0.0309)** | **** (0.0021)** |
| | 4C bloodstream | ns (0.1884) | **** (0.0025)** |
| | 2C oral/vag | ns (0.9492) | ***** (0.0003)** |
| | 4C oral/vag | ns (0.1728) | ns (0.0671) |
| 4C Lab hom | 2C Lab het | ****** (<0.0001)** | ns (0.1135) |
| | 4C lab het | ****** (<0.0001)** | ns (0.1660) |
| | 2C bloodstream | ****** (<0.0001)** | ns (0.1362) |
| | 4C bloodstream | ns (0.7349) | ns (0.2955) |
| | 2C oral/vag | ns (0.1165) | ns (0.1023) |
| | 4C oral/vag | ns (0.7460) | ns (0.8598) |
| 2C lab het | 4C lab het | ns (0.2505) | ns (0.7200) |
| | 2C bloodstream | ns (0.9813) | ns (0.7254) |
| | 4C bloodstream | ***** (0.0002)** | ns (0.6838) |
| | 2C oral/vag | *** (0.0267)** | ns (0.2165) |
| | 4C oral/vag | ***** (0.0002)** | ns (0.1229) |
| 4C lab het | 2C bloodstream | ns (9.3855) | ns (0.9048) |
| | 4C bloodstream | ***** (0.0005)** | ns (0.5095) |
| | 2C oral/vag | **** (0.0034)** | ns (0.5684) |
| | 4C oral/vag | ***** (0.0002)** | ns(0.0953) |
| 2C bloodstream | 4C bloodstream | ***** (0.0006)** | ns (0.4838) |
| | 2C oral/vag | *** (0.0486)** | ns (0.4857) |
| | 4C oral/vag | **** (0.0013)** | ns (0.1824) |
| 4C bloodstream | 2C oral/vag | ns (0.3885) | ns (0.1408) |
| | 4C oral/vag | ns (0.9500) | ns (0.2732) |
| 2C oral/vag | 4C oral/vag | ns (0.2538) | *** (0.0224)** |

Table S5: Pairwise reproductive timing comparisons (Mann Whitney test) for uninfected and all eight *C. albicans* strains, for each host genotype.
